# Corticosteroids prevent pathological angiogenesis yet compromise reparative vascular remodeling in the retina

**DOI:** 10.1101/2024.06.06.597805

**Authors:** Masayuki Hata, Maki Hata, Agnieszka Dejda, Frédérique Pilon, Roberto Diaz, Frédérik Fournier, Jean-Sebastien Joyal, Gael Cagnone, Yotaro Ochi, Sergio Crespo-Garcia, Ariel Wilson, Przemyslaw Sapieha

**Affiliations:** Departments of Ophthalmology, University of Montreal, Montreal, Quebec, H1T 2M4, Canada; Departments of Biochemistry and Molecular Medicine, Maisonneuve-Rosemont Hospital Research Centre, University of Montreal, Montreal, Quebec, H1T 2M4, Canada; Departments of Pediatrics, Ophthalmology, and Pharmacology, Centre Hospitalier Universitaire Ste-Justine Research Center, Montréal (Quebec), H3T 1C5, Canada; Department of Pathology and Tumour Biology, Graduate School of Medicine, Kyoto University, Kyoto, Japan

**Keywords:** diabetic retinopathy, corticosteroids, dexamethasone, angiogenesis, inflammation, vascular remodeling

## Abstract

Tissue inflammation is often broadly associated with cellular damage, yet sterile inflammation also plays critical roles in beneficial tissue remodeling. In the central nervous system (CNS), this is observed through a predominantly innate immune response in retinal vascular diseases such as age-related macular degeneration, diabetic retinopathy and retinopathy of prematurity. Here we set out to elucidate the dynamics of the immune response during progression and regression of pathological neovascularization in retinopathy. In a mouse model of oxygen-induced retinopathy, we report that broad spectrum corticosteroid drugs such as dexamethasone suppress initial formation of pathological pre-retinal neovascularization in early stages of disease, yet blunt successive waves of reparative inflammation and hence prevent beneficial vascular remodeling. Using genetic depletion of distinct components of the innate immune response, we demonstrate that CX3C chemokine receptor 1 (CX3CR1)-expressing microglia contribute to angiogenesis. Conversely, myeloid cells expressing Lyz-M (lysozyme 2) are recruited to sites of damaged blood vessels and pathological neovascularization where they partake in a reparative process that ultimately restores circulatory homeostasis to the retina. Hence, the Janus-faced properties of anti-inflammatory drugs should be considered when treating retinal vascular disease and particularly in stages associated with persistent neovascularization.

## INTRODUCTION

Retinal homeostasis is reliant on innate immunity^1^. Microglia, monocytes, mononuclear phagocytes (MNPs) and neutrophils have been well studied in the sterile inflammatory response during retinal disease^2,3^ where they partake in mediating both seemingly destructive processes such as pathological angiogenesis^4,5,6^ and phagocytosis of damaged photoreceptor cells^7,8^ as well as reparative events such as pruning diseased blood vessels^9,3^. The dual function of the immune response suggests that use of anti-inflammatory strategies must be staged in order to preserve beneficial properties of the immune response while blunting excessive cytokine production in order to ensure retinal homeostasis and proper sight.

Corticosteroid and specifically glucocorticoid drugs are amongst the most widely used pharmaceutical products in the industrialized world due to their potent immunomodulatory effects^10,11^. The first clinical confirmation for corticosteroids came in the 1930s when animal adrenocortical tissue was shown to improve outcome of adrenal failure in humans^12^ and first patients were treated with cortisone for rheumatoid arthritis in 1948^13^. Chemical modifications of the steroid skeleton led to highly potent anti-inflammatory glucocorticoid such as dexamethasone that have been in use since the late 1950s^14^. In ophthalmology, corticosteroids are currently widely used for a variety of ophthalmic conditions with local administration for conditions such as conjunctivitis and keratitis, non-infectious posterior uveitis, macular edema due to retinal vasculitis and diabetic retinopathy and systemic administration for scleritis and optic neuritis^15,16^. Given the broad-spectrum of action of corticosteroids, their indication typically implies a lack of clear mechanistic understanding of underlying disease process.

In light of common use of corticosteroids in clinical treatment of prevalent sight threatening complications such as diabetic macular edema^17,18^, we explored the consequence of intravitreal dexamethasone treatment on tissue remodeling in a mouse model of pathological retinal neovascularization^19^. We focused on distinct stages of retinopathy such as pathological neovascularization and beneficial vascular remodeling. Additionally, we investigate the contribution of local CX3CR1^+^ resident immune cells and myeloid-derived LyzM^+^ retinal mononuclear phagocytes in vascular remodeling during retinopathy.

## RESULTS

### Dexamethasone treatment suppresses pathological neovascularization in early stages but prevents reparative vascular remodeling in later stages of disease

To investigate the effects of anti-inflammatory drugs such as glucocorticoid steroids on pathological vascularization in the retina, we employed the mouse model of oxygen-induced retinopathy (OIR) that is characterized by ischemic retinal tissues and deregulated angiogenesis^19^. Mouse pups were exposed to 75% oxygen from postnatal day (P) 7 to P12 to trigger vaso-obliteration, then returned to room air to initiate a second phase of pathological neovascularization that peaks at P17 and is followed by a phase of vascular regression (**Fig 1A).** To elucidate general biological processes at play during the progression of retinopathy, we conducted a time-course analysis of transciptomic changes through bulk-RNA sequencing during distinct stages of OIR^9,20,21^ at P14 (during neovascularization), P17 (at maximal neovascularization), and P30 (following vascular normalization) and investigated enriched gene sets using GO (Gene Ontology). We observed transcript enrichment in processes related to inflammation/immune response, metabolic process, angiogenesis, and hypoxia at P14 during initiation of prereitnal neovascularization, and inflammation/immune response, wound healing, and angiogenesis at P17 of OIR during peak neovascularization with **(Fig 1B and Supplemental Fig 1**). At P30, following vascular normalization, inflammatory genes are regulated to a lesser extent and genes corresponding to „*negative regulation of immune system process*“ apear to be higly regulated.

**Figure 1.**
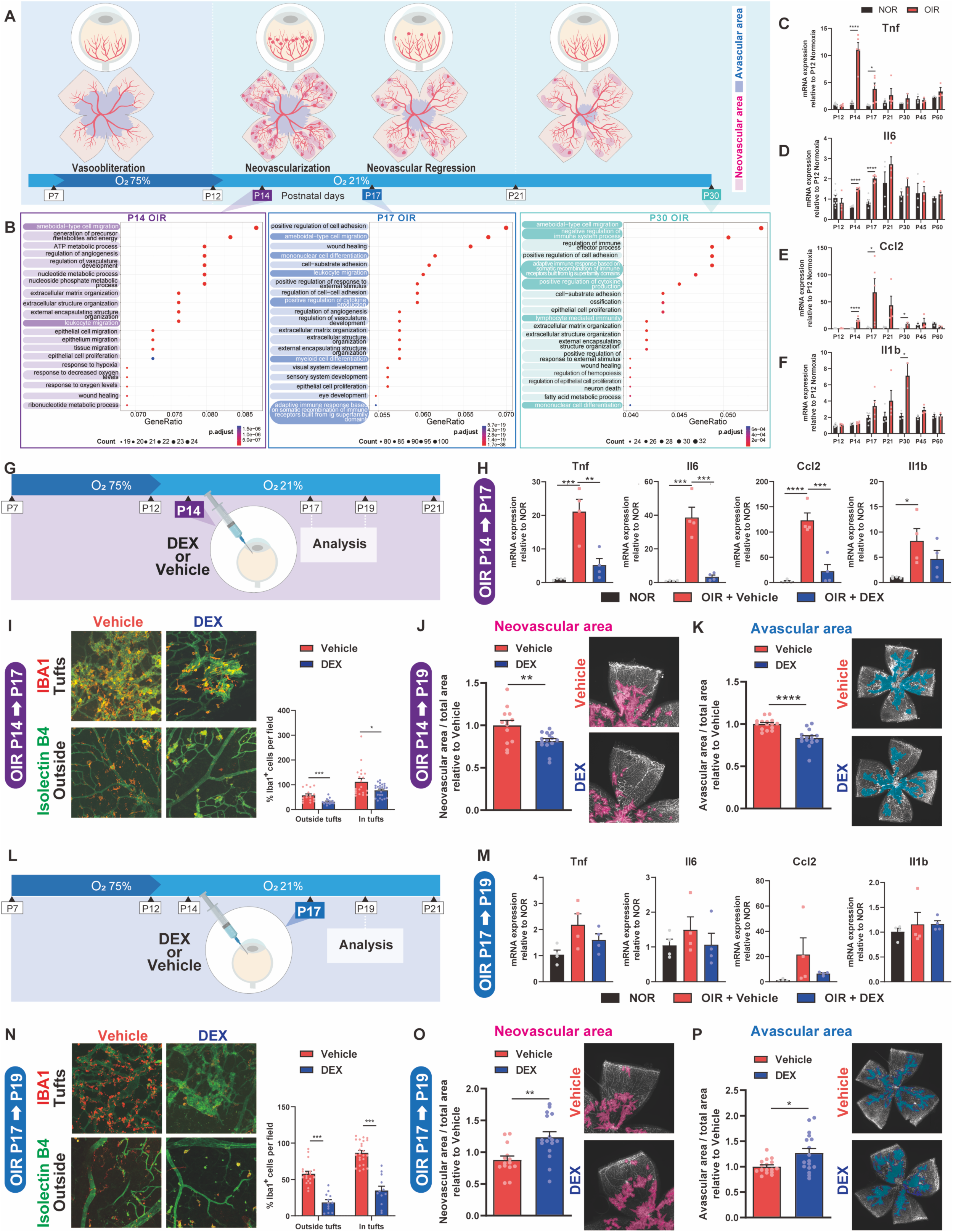
Intravitreal Dexamethasone treatment suppresses pathological neovascularization in early stages but prevents reparative vascular remodeling in later stages of disease. (A) Schematic representation of the mouse model of OIR and the distinct phases of the pathological vascularization (vasoobliteration from P7 to P12, neovascularization from P12 to P17, and neovascular regression from P17). (B) Gene ontologies (GO) related to the biological processes for bulk RNA-seq from OIR and normoxic mouse retinas at P14, P17, and P30 (N = 2 to 3 mice per condition). Inflammation-related GO terms are highlighted. (C-F) mRNA expression of inflammation-related genes (C) *Tnf,* (D) *Il6,* (E) *Ccl2,* and (F) *Il1b* throughout the progression of OIR. Data are presented as fold change compared with P12 normoxic retinas (N = 3 to 10 depending on the group). Statistics were calculated comparing OIR versus normoxia for each given time point. (G) Schematic representation of intravitreal administration of dexamethasone (0.3mg) to P14 pups during OIR. (H) mRNA expression of *Tnf, Il6*, *Ccl2,* and *Il1b* (N = 4 per condition) in P17 retinas of OIR treated with or without dexamethasone (OIR + DEX, OIR + Vehicle, respectively) relative to normoxia. (I) IBA1^+^ phagocytes colocalized with isolectin-B4^+^ ECs were found in P17 flatmounts of OIR treated with or without dexamethasone (DEX and Vehicle, respectively) in neovascular tuft areas and outside tuft areas (N = 16 to 24 depending on the group). (J and K) Neovascular area (J) and avascular area (K) as assessed at P19 flatmounts of OIR treated with or without dexamethasone (N = 13 to 15 depending on the group). (L) Schematic representation of intravitreal administration of dexamethasone (0.3mg) to P17 pups during OIR. (M) mRNA expression of *Tnf, Il6*, *Ccl2,* and *Il1b* (N = 4 per condition) in P19 retinas of OIR treated with or without dexamethasone (OIR + DEX, OIR + Vehicle, respectively) relative to normoxia. (N) IBA1^+^ phagocytes colocalized with isolectin-B4^+^ ECs were found in P19 flatmounts of OIR treated with or without dexamethasone (DEX and Vehicle, respectively) in neovascular tuft areas and outside tuft areas (N = 12 to 24 depending on the group). (O and P) Neovascular area (O) and avascular area (P) as assessed at P19 flatmounts of OIR treated with or without dexamethasone (N = 14 to 17 depending on the group). Student’s unpaired t-test (C-F, I-K, N-P) and one-way ANOVA with Tukey’s multiple-comparison test (H, M) were used; *P < 0.05, **P < 0.01, ***P < 0.001 ****P< 0.0001; error bars represent mean ± SEM.

To verify these findings, we investigated individual transcripts by quantitative PCR (qPCR). We observed induction of *Tnf*, *Il6,* and *Ccl2* throughout the course of pathological neovascularization (P13-P17) and vascular regression (P17-P21) (**Fig 1C-F**). *Il1b* was induced around peak neovascularization and persisted (**Fig 1F).**

Next, we evaluated the effects of anti-inflammatory treatments on tissue repair, (specifically vascular remodeling) during retinopathy. We injected 0.3mg of dexamethasone into the vitreous of C57BL6 mice at P14, during onset of neovascularization (**Fig 1G**). As expected, at P17, dexamethasone suppressed expression of *Tnf*, *Il6*, *Il1b*, and *Ccl2* as assessed by qPCR (**Fig 1H**). Accordingly, dexamethasone-treated eyes had fewer IBA1-expressing phagocytes, both within and outside neovascular tufts (**Fig 1I)** measured in retinal flatmounts. Importantly, compared to vehicle controls, treatment of OIR retinas with dexamethasone (DEX) at P14 significantly reduced areas of pathological neovascularization (**Fig 1J)** and increased vascular regeneration (**Fig 1K**) as assessed at P19.

To investigate the impact of intravitreal dexamethasone on vascular remodeling, we next treated mouse pups at P17 during maximal neovascularization (**Fig 1L**). Evaluation of retinas at P19 revealed that dexamethasone led to a non-significant trend in decrease of *Tnf*, *Il6*, and *Ccl2* (**Fig 1M**) accompanied by a decrease in IBA1-positive inflammatory phagocytes both inside and outside of neovascular tufts (**Fig 1N**). In contrast to early treatment with dexamethasone at P14, late administration at P17 significantly compromised regression of pathological neovascularization and slowed vascular regeneration (**Fig 1O and 1P**). Collectively, intravitreal dexamethasone in early stages of neovascularization prevented pathological angiogenesis, yet later administration blunted beneficial vascular remodeling.

### Intravitreal Dexamethasone reduces the number of endothelial cells undergoing apoptosis within pathological neovascularization

Pruning of pathological neovascularization requires elimination of endothelial cells through apoptotic mechanisms^9,3^. Using flowcytometry during stages of vascular normalization at P19 OIR, we found increased numbers of Annexin V-positive endothelial cells during regression of pathological blood vessels (**Supplemental Fig 2A, B).** Consistent with compromised vascular remodeling, retinas from mice treated with dexamethasone had significantly fewer Annexin V-positive endothelial cells (**Supplemental Fig 2A, B)**. Cleaved caspase3 staining confirmed that apoptotic cells were primarily confined to sites of pathological neovascularization (tufts) (**Supplemental Fig 2C, D)** and quantification confirmed that dexamethasone significantly reduced cleaved caspase 3-expressing apoptotic cells within pathological neovascularization. Altogether, this data is consistent with compromised clearance of pathological blood vessels following intravitreal use of dexamethasone.

### Distinct innate immune cell populations contribute to neovascularization and vascular remodeling of OIR

Corticosteroids such as dexamethasone, have broad immunosuppressive effects, independent of immune cell population. Consistent with this, flowcytometry analysis showed dexamethasone administration decreased lymphocytes (CD45^+^/CD11b^-^/CD3ɛ^+^), microglia (CD45^+^/CD11b^+^/CX3CR1^+^), and MNPs (CD45^+^/CD11b^+^/Ly6C^int/high^/Ly6G^low^) in the retina at P17 of OIR (**Fig 2A, with gating in Supplemental Fig 3)**, and microglia, MNPs, and neutrophils (CD45^+^/CD11b^+^/Ly6C^int^/Ly6G^high^) in the retina at P19 of OIR (**Fig 2B**).

**Figure 2.**
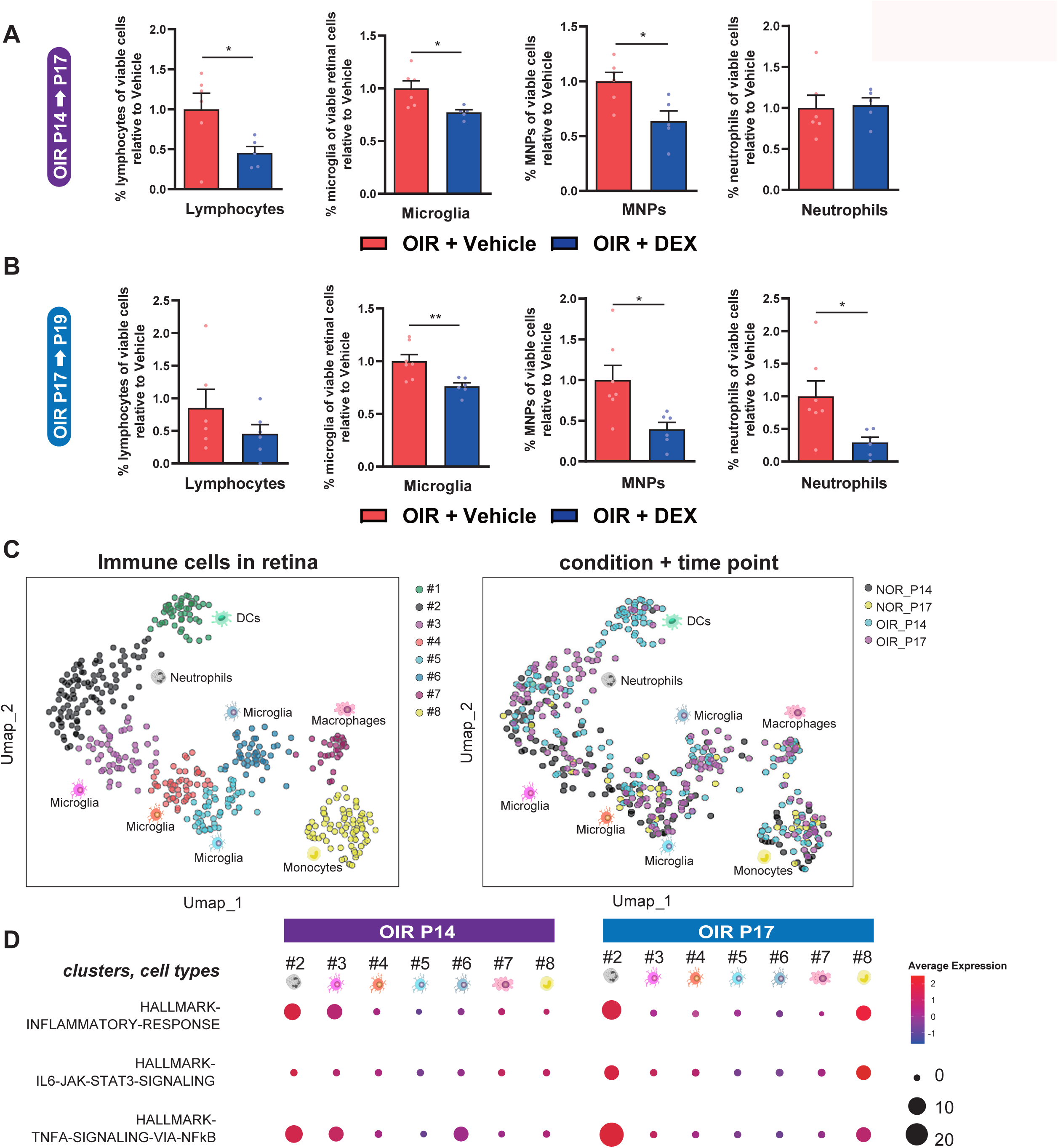
Distinct populations of innate immune cells are elevated during neovascularization and vascular remodeling in ischemic retinopathy. **(A and B)** Flow cytometric analyses of OIR retinas treated with and without dexamethasone (OIR+DEX and OIR+Vehicle, respectively). Percentages of viable lymphocytes (CD45^+^/CD11b^-^/CD3ɛ^+^), microglia (CD45^+^/CD11b^+^/CX3CR1^+^), mononuclear phagocytes (MNPs; CD45^+^/CD11b^+^/Ly6C^int/high^/Ly6G^low^), or neutrophils (CD45^+^/CD11b^+^/Ly6C^int^/Ly6G^high^) assessed at P17 **(A)** and P19 **(B)** by FACS (N = 5 to 7 depending on the group). **(C)** Uniform manifold approximation and projection (UMAP) plot for single-cell RNAseq of retinal immune cells from normoxic and OIR retinas at P14 and P17 (from two normoxic and three OIR sets of data). UMAP split according to immune cells populations (left panel) and condition+timepoint (right panel). Each dot represents one cell. **(D)** Dot plot representing results of GSVA for the inflammation-associated gene sets from Hallmark in each of the seven populations identified in (**C**) at P14 and P17 of OIR. Student’s unpaired t-test (**A, B**) were used; *P < 0.05, **P < 0.01; error bars represent mean ± SEM.

To identify potential cell populations associated with clearance of pathological neovascularization from peak neovascularization at P17 of OIR, we performed scRNA sequencing (scRNA-seq). Principal components analysis (PCA) and uniform manifold approximation and projection (UMAP) plot of different clustered retinal immune cell types with similar transcriptional profiles revealed 8 independent subpopulations present in the retina (**Fig 2C**). Marker gene expression analysis identified cluster #1 as Dendritic Cells (DCs), #2 as neutrophils, #3-6 as microglia, #7 as macrophages, and #8 as monocytes (**Supplemental Fig 4**). Consistent with qPCR data above (**Fig 1C-F**), pathway analysis confirmed increased Tnf signaling at P14, whereas IL6 signaling was induced at P17 (**Fig 2D**). Notably, cluster #2 (neutrophils) and #3 (microglia) showed the most enrichment in inflammatory pathways at P14, while #2 (neutrophils) and #8 (monocytes) showed the most enrichment in inflammatory processes at P17.

### LysM^+^ monocytes mediate vascular remodeling in later stages of OIR while CX3CR1-expressing cells primarily contributing to angiogenesis

scRNA-seq data revealed monocytes (cluster #8) expressed LyzM but not CX3CR1, while CX3CR1 was highly expressed in microglia (cluster #4) and to a lesser extent in monocytes (cluster #8) (**Fig 3A**). To assess the contribution of each population of innate immune cells on pathological angiogenesis and vascular remodeling, we generated mice expressing diphtheria toxin (DTx) receptor under control of LyzM or CX3CR1 promoters. These mice allow investigation of the contribution of each cell population to vascular phenotypes through selective and time-specific ablation of monocyte-derived LyzM-expressing cells or microglia-expressing CX3CR1 secondary to intravitreal administration of DTx ^22^.

**Figure 3.**
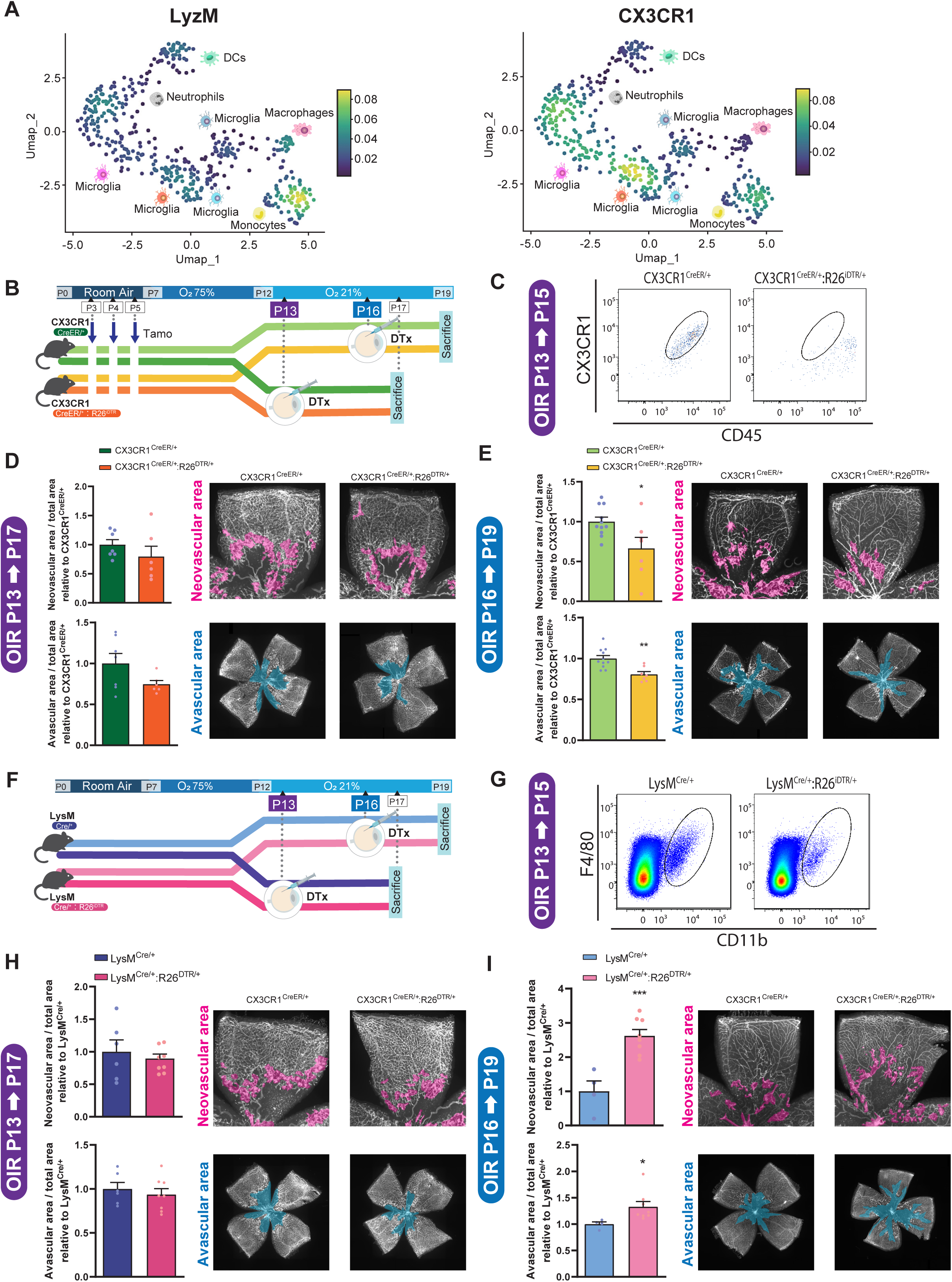
LysM^+^ monocytes mediate vascular remodeling in later stages of OIR while CX3CR1-expressing cells contribute to angiogenesis. (**A**) UMAP visualization of the expression of two major myeloid marker genes LysM (left panel) and CX3CR1 (right panel) in retinal immune cells during OIR. (**B**) Time course of *Cx3cr1*^CreER/+^ and *Cx3cr1* ^CreER/+^:R26^iDTR/+^ OIR mice. For both *Cx3cr1* ^CreER/+^ and *Cx3cr1* ^CreER/+^:R26 ^iDTR/+^ mice, tamoxifen (TAM) was administered daily between P3 and P5, and diphtheria toxin intravitreally (ivt) at either P13 or P16. (**C**) Representative FACS plots of CX3CR1^+^ microglia of retinas from CX3CR1 ^CreER/+^ and CX3CR1 ^CreER/+^ :R26 ^iDTR/+^ mice following intravitreal diphtheria toxin injection. (**D**) Neovascular area and avascular area as assessed at P17 flatmounts of OIR treated with or without dexamethasone at P13 (N = 6 or 7 depending on the group). (**E**) Neovascular area and avascular area as assessed at P19 flatmounts of OIR treated with or without dexamethasone at P16 (N = 7 or 11 depending on the group). (**F**) Time course of *LysM*^Cre/+^ and *LysM*^Cre/+^:R26^iDTR/+^ OIR mice. For both *LysM*^Cre/+^ and *LysM*^Cre/+^:R26^iDTR/+^ mice, tamoxifen (TAM) was administered daily between P3 and P5, and diphtheria toxin intravitreally (ivt) at either P13 or P16. (**G**) Representative FACS plots of MNPs of retinas from *LysM*^Cre/+^ and *LysM*^Cre/+^:R26^iDTR/+^ mice following intravitreal diphtheria toxin injection. (**H**) Neovascular area and avascular area as assessed at P17 flatmounts of OIR treated with or without dexamethasone at P13 (N = 6 or 8 depending on the group). (**I**) Neovascular area and avascular area as assessed at P19 flatmounts of OIR treated with or without dexamethasone at P16 (N = 4 or 8 depending on the group). Student’s unpaired t-test (**D, E, H, I**) were used; *P < 0.05, **P < 0.01, ***P < 0.001; error bars represent mean ± SEM.

When CX3CR1-positive cells were eliminated during the phase of pathological angiogenesis at P13 (**Fig 3B and 3C)** a slight trend to lower pathological neovascularization vascular regeneration at P17 was observed (**Fig 3D**). In contrast, when CX3CR1-positive cells were removed at P16 towards peak pathological angiogenesis and onset of vascular remodeling, we observed a significant decrease in both areas of pathological neovascularization and extent of vascular regeneration at P19 (**Fig 3E**).

Next, we eliminated LysM-positive cells during neovascularization (P13) and vascular remodeling (P16) with intravitreal administration of DTx **(Fig 3F** and 3**G)**. Similar to CX3CR1-positive cells, depletion of LysM-positive cells at P13 lead to a non-significant trend towards lower pathological neovascularization nor vascular regeneration at P17 (**Fig 3H**). However, removal of LysM-positive cells at P16 significantly compromised beneficial vascular remodeling and led to persistent pathological neovascularization and stalled vascular remodeling to a similar extent as what we observed with intravitreal dexamethasone (**Fig 3I**). In agreement with scRNAseq data showing that neutrophils and monocytes upregulate most robustly their inflammatory profiles prior to vascular regression (**Fig 2D**), these results indicate that CX3CR1-positive and LysM-positive myeloid cells play distinct roles in vascular remodeling, with LysM-positive myeloid cells partaking in the clearance of pathological neovascularization and CX3CR1-expressing cells contributing to pathological angiogenesis.

## DISCUSSION

While non-resolving inflammation is typically associated with tissue damage and disease, the innate immune system’s primary function is to fend off non-self pathogens and ensure and tissue repair and homeostasis ^23,24^. In this study we put forward two sets of data highlighting that non-specific inhibition of effectors of retinal immunity with corticosteroids can inhibit formation of pathological angiogenesis, yet compromise intrinsic repair mechanisms that themselves remodel and eliminate diseased neovascularization. In addition, we provide evidence for a role for monocyte-like cells in driving beneficial vascular remodeling and microglia in mediating retinal neo-angiogenesis.

Our data are in line with the notion that corticosteroids prevent angiogenesis associated with disease, and current clinical data that demonstrate topical dexamethasone can reduce macular edema^25^ and progression to severe Type II neovascular ROP^26^. Yet, our data also highlight that the retinal immune system is critical for reparative vascular remodeling and non-specifically dampening the inflammatory response at disease stages when vascular remodeling occurs can compromise tissue repair. Using mouse genetics, we demonstrate that the reparative properties of the innate immune system can be attributed LyzM-expressing cells that are likely of monocytic origin. These finding agree with our previous work demonstrating enrichment of senescent endothelial cells in neovascular tufts^21,27^ with distinct secretomes that recruit effectors of innate immunity such as neutrophils^9^. Neutrophils in turn extrude DNA and form Neutrophil Extracellular Traps (NETs) that are strands of decondensed DNA ^28^. NETs target senescent blood vessels for vascular remodeling, and depleting neutrophils or inhibiting NETs reduces vascular remodeling in retinopathy^28^ to a similar extent as what we observed with dexamethasone treatment. Interestingly, our data suggest that intravitreal dexamethasone reduces numbers of neutrophils and specifically in later stages of OIR associated with peak neovascularization. Given that neutrophils express LyzM, their elimination from _LysM_Cre/+_:R26_DTR/+ may be part of the mechanism to prevent regression of pathological angiogenesis. Similarly, mice that are deficient in monocyte-chemoattractant protein-1 (MCP-1) have delayed regression of neovascular tufts^3^, further supporting the importance proper innate immune function in retinal vascular homeostasis.

Collectively, our data support use of broad-spectrum anti-inflammatory drugs in treatment of active neovascular retinal disease, yet suggest that their use should be disease-stage specific in order to avoid compromising endogenous repair mechanisms of retina and other tissues. While initial triggers of pathological angiogenesis may be governed by components of the innate immune system, impairing inflammation at the height of neovascularization in late stages of retinopathy may compromise beneficial vascular remodeling^9^.

## METHODS

### Mice

All animal studies were performed in compliance with the ARRIVE guidelines and the Association for Research in Vision and Ophthalmology (ARVO) Statement for the Use of Animals in Ophthalmic and Vision Research and were approved by the Animal Care Committee of the Maisonneuve-Rosemont Hospital Research Center in agreement with the guidelines established by the Canadian Council on Animal Care.

Animals were housed in the animal facility of the Hospital Maisonneuve-Rosemont Research Center under a 12h light/dark cycle with *ad libitum* access to food and water unless indicated otherwise.

Homozygous *B6.129P2(C)-Cx3cr1^tm2.1(cre/ERT2)Jung^/J* (referred as *CX3CR1^CreER^*) mice were crossed in-house with homozygous *C57BL/6-Gt(ROSA)26Sor^tm1(HBEGF)Awai^/J* (referred as *R26^iDTR^*) mice to obtain heterozygous *CX3CR1^CreER/+^:R26 ^iDTR/+^* mice.

Homozygous B6.129P2-*Lyz2^tm1(cre)Ifo^*/J (referred to as LysMcre) mice were crossed with homozygous *C57BL/6-Gt(ROSA)26Sor^tm1(HBEGF)Awai^/J* (referred as *R26^iDTR^*) mice to obtain heterozygous or *LysM^Cre/+^:R26 ^iDTR/+^* mice.

### Oxygen-induced retinopathy mouse model

Oxygen-induced retinopathy (OIR) was carried out as described previously ^29^. In brief, mouse pups (C57Bl/6J, *CX3CR1^CreER/+^:R26 ^iDTR/+^* or *LysM^Cre/+^:R26 ^iDTR/+^*mice) and their corresponding fostering mothers (CD1 female) were exposed to 75% O_2_ from P7 to P12 and returned to room air afterwards. This model resembles aggressive neovascular features as in human ocular neovascular diseases such as proliferative DR ^19^ ^30^. Upon return to room air, hypoxia-driven epi-retinal neovascularization develops from P14 onward ^31^.

### Myeloid cell depletion

Microglia depletion was performed using CX3CR1CreER/+:R26 iDTR/+ in which the activation of Cre recombinase (under the control of the Cx3cr1 promoter) can be induced by tamoxifen treatment and leads to surface expression of DTR on CX3CR1-expressing cells. At P3, P4 and P5, mice were subjected to daily intraperitoneal injections with tamoxifen diluted in corn oil (0.05 mg per mouse per day) for three consecutive days.

To deplete CX3CR1^+^ cells in *CX3CR1^CreER/+^:R26 ^iDTR/+^* mice, diphtheria toxin was administered intravitreally (25 ng/1 μl saline per eye) at P13 or P16. Depletion of LysM^+^ cells in *LysM^Cre/+^:R26 ^iDTR/+^* mice was achieved similarly at P13 and P16.

### Gene set enrichment analysis from mouse retina RNA-Seq

GSEA was conducted using GSEA v4.0.1 software provided by Broad Institute of Massachusetts Institute of Technology and Harvard University. We used GSEA to validate correlation between molecular signatures in phenotypes of interest. Enrichment analysis was conducted on pre-ranked lists based on shrunken log2 fold changes from DESeq2 lfcShrink option. Default parameters were changed as follows: Gene sets of interest were found in a catalog of functional annotated gene sets from MSigDB; phenotype label was defined as “OIR” versus “Normoxia”; gene sets smaller than 15 and larger than 500 were excluded from the analysis; statistic used to score hits was defined as “weighted p1”.

### Bulk RNA-seq analysis

Bulk RNA-seq data was described in Ref^20, 21^. edgeR package in R was used to identify the differentially expressed genes in OIR compared to normoxia at each time point, with an FDR threshold of 0.05. The analysis was performed in genes expressed at >1 count per million in two or more samples, and generalized linear models were used to compare gene expression data. Gene ontology (GO) analysis was performed using significantly up- or down-regulated genes and the enrichGO function from the clusterProfiler package with biological process ontologies, and top 20 genes ordered by gene ratio were plotted.

### Droplet-based single-cell RNA sequencing analysis

Unique molecular identifier (UMI) counts for normoxic and OIR retina scRNA-Seq replicates were merged into one single Digital Gene Expression (DGE) matrix and processed using the Seurat package (Spatial reconstruction of single-cell gene expression data). Cells expressing less than 100 genes and more than 10% of mitochondrial genes were filtered out. Single cell transcriptomes were normalized by dividing by the total number of UMIs per cell, then multiplying by 10,000. All calculations and data were then performed in log space (i.e. ln(transcripts-per-10,000 +1)). After aligning whole and rod-depleted datasets using canonical correlation analysis on the most variable genes in the DGE matrix, PCA analysis identified 20 significant PC which served as input for dimensionality reduction and embedding into 2-dimentional space. To identify putative cell types in the embedded space, we used a density clustering approach and computed average gene expression for each of the identified cluster based on Euclidean distances. We then compared each of the different clusters to identify marker genes that were significantly enriched for each cluster. Transcriptomic differences between normoxic and OIR cell types were statistically compared using a negative binomial model and analyzed using visualization tools including RidgePlot and UMAP plot from the Seurat R Package. Single-cell gene expression profiles from each separate cell type identified by scRNA-Seq were further analyzed using Gene Set Variation Analysis (GSVA).

### Intravitreal injections

Pups were injected intravitreally at P14 using a Hamilton syringe (10µl) with a glass capillary coupled onto it. Each treated pup had both eyes injected with either 1µl dexamethasone (0.3mg) of 1µl of appropriate vehicle control. Animals were evaluated at different time-points depending on the experiment.

### Transcription analysis by Quantitative PCR

RNA extraction was performed with snap frozen mouse retinas using TRIzol reagent (Cat# 15596026; Invitrogen) and digested with DNase I (Sigma Aldrich; Cat# D4527) following manufacturer instructions to avoid genomic DNA amplification. Total RNA was reverse transcribed using a 5X All-In-One RT MasterMix (Cat# G590; Applied Biological Materials Inc.) according to the manufacturer’s instructions. Gene expression was analyzed using Bright Green2X qPCR Master Mix-Low Rox in an Applied Biosystems 7500 Real-Time PCR System (Thermo Fisher Scientific, Waltham, MA, USA). Primer sequences used in this study are listed in the Supplemental Table 1. Analysis of expression was followed using the ΔΔCT method. Actb expression was used as the reference housekeeping gene. Statistical analysis was performed on ΔΔCT values, and data was represented as the expression of the target genes normalized to Actb (folds of increase).

### Retinal immunohistochemistry

Enucleated eyes were fixed in PFA 4% during 1 hour at RT and washed with PBS. Retina was micro-dissected and incubated in a blocking solution containing 3% of bovine serum albumin (BSA) during 1 hour. Antibodies including fluorescence-labeled ISOLECTIN-B_4_ (Cat# FL-1101, Vector Labs) and Cleaved Caspase-3 (Cat# 9661S, Cell Signaling Technologies) were diluted using the same solution and applied on the tissue overnight at 4°C. After washes with PBS, species-appropriate fluorescence-conjugated secondary antibodies were applied for 1 hour at RT. Samples were counterstained with DAPI, mounted and imaged using a Olympus FV1000 confocal microscope (Olympus Canada, Richmond Hill, ON, Canada).

### Analysis of retinal vasculature

Enucleated eyes were fixed in PFA 4% for 1 hour at RT. Retinas were carefully dissected and incubated overnight at 4°C with fluorescence-labeled ISOLECTIN-B_4_ (IB_4_) in PBS. After mounting, retinas were imaged using a Zeiss AxioObserver.Z1 microscope (Zeiss, Jena, Germany). To determine extent of avascular area or neovascularization area, images were processed using ImageJ (U. S. National Institutes of Health, Bethesda, Maryland, USA) and analyzed by the SWIFT-neovascularization method as described previously^31^. Results are expressed normalized to control.

### Fluorescence-Activated Cell Sorting (FACS) on retina and RPE-choroid-sclera complexes

Retinas and RPE-choroid-sclera complexes were cut into small pieces and homogenized in a solution of 750U/mL DNAse I (Cat# D4527, Sigma-Aldrich) and 0.5mg/mL of collagenase D (Cat# 11088866001, Roche) for 20 minutes at 37°C. Homogenates were filtered through a 70 μ m cell strainer, counted and resuspended in PBS +3% FBS. Viability of the cells was checked by 7-ADD (Cat# 420404: Biolegend) staining for 20 min at room temperature. After incubation with LEAF purified anti-mouse CD16/32 (Cat# 101310; BioLegend) for 10 minutes at 4°C to block FC receptors, cells were incubated for 25 minutes at 4°C with the following antibodies: BV711 anti-mouse/human CD11b (Cat# 101242; BioLegend), PE anti-mouse CX3CR1 (Cat# FAB5825P; R&D), APC anti-mouse CD45.2 (Cat# 109814; BioLegend), APC/Cy7 anti-mouse Ly-6G (Cat# 127624; BioLegend), and PE/Cy7 anti-mouse F4/80 (Cat# 123114; BioLegend), or BV785 anti-mouse CD45.2 (Cat# 109839; BioLegend), FITC anti-mouse CD31 (Cat# 102506; BioLegend), and APC Annexin-V (Cat# 640919; BioLegend). Fluorescence-activated cell sorting (FACS) was performed on a BD LSR FortessaTM X-20 cell analyzer, and data were analyzed using FlowJo software (Version 10.2; FlowJo, Ashland, OR, USA).

### Statistical analyses

Data are expressed as mean ± SEM, unless indicated otherwise. Multiple Student’s t-tests, 2-tailed Student’s *t* test or ANOVA t-test were used to compare two or more than two groups, respectively. Statistical significance was considered when *p*<0.05 and noted as: * *p*<0.05, ** *p*<0.01, *** *p*<0.001 and **** *p*<0.0001. All experimental Ns are indicated in the figure legends.

## Data and code availability

RNAseq and Single-cell RNA-seq data are deposited in NCBI’s Gene Expression Omnibus and are accessible through GEO accession numbers GSE158799 and GSE150703.

