## Supplemental figures and tables for "Corticosteroids prevent pathological angiogenesis yet compromise reparative vascular remodeling in the retina"

**Title:**

**Affiliations:**

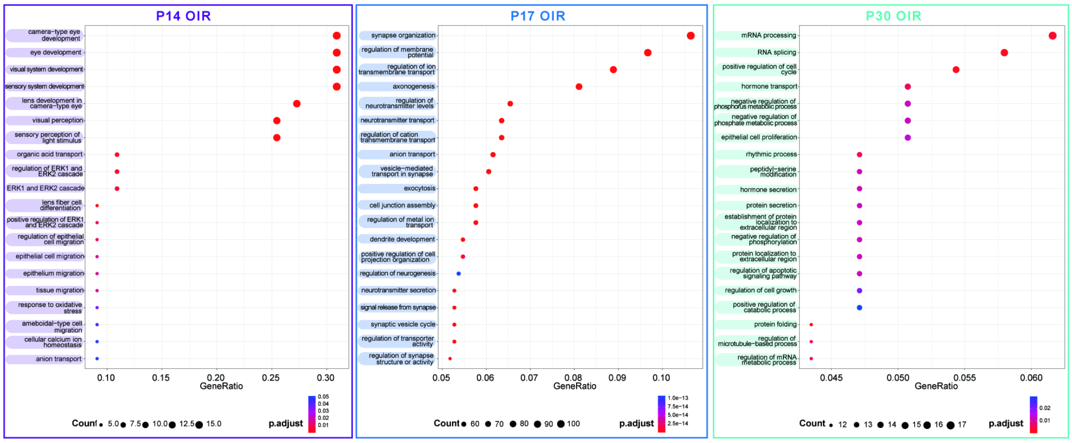


**Fig. S1.** Down-regulated gene ontologies (GO) related to the biological processes for bulk RNA-seq from OIR and normoxic mouse retinas at P14, P17, and P30 (n = 2 to 3 mice per condition).

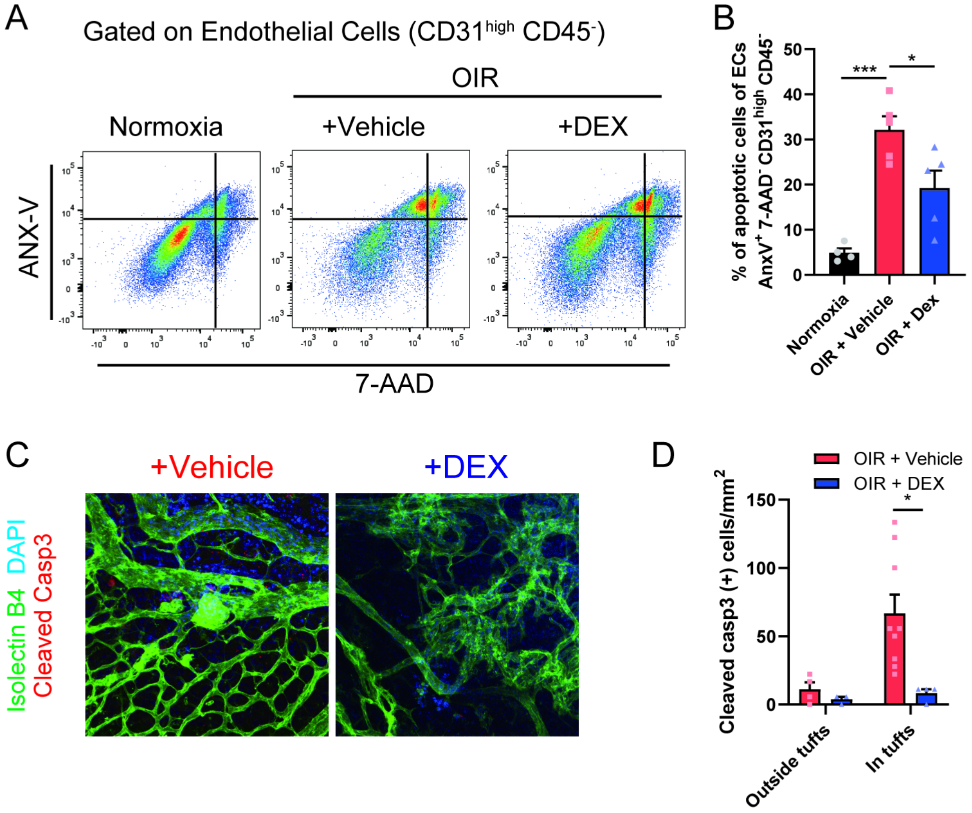


**Fig. S2.** Dexamethasone treatment suppress apoptosis of endothelial cells within pathological vasculature during vascular remodeling. (**A and B**) Flow cytometric analyses of AnexinV-positive endothelial cells of OIR retinas treated with and without dexamethasone (OIR+DEX and OIR+Vehicle, respectively). Percentages of apoptotic cells ($AnxV^+$/7-$AAD^-$/$CD31^{high}$/$CD45^-$) and dead cells (7-$AAD^+$/$CD31^{high}$/$CD45^-$) of endothelial cells assessed at P19 by FACS (N = 4 to 5 depending on the group). (**C and D**) Cleaved caspase-$3^+$ apoptotic cells colocalized with isolectin-$B4^+$ ECs were found in P19 flatmounts of OIR treated with or without dexamethasone (DEX and Vehicle, respectively) in neovascular tuft areas and outside tuft areas (N = 3 to 6 depending on the group).

One-way ANOVA with Tukey's multiple-comparison test (**B**) and student's unpaired t-test (**D**) were used; *$P < 0.05$, ***$P < 0.001$; error bars represent mean ± SEM.

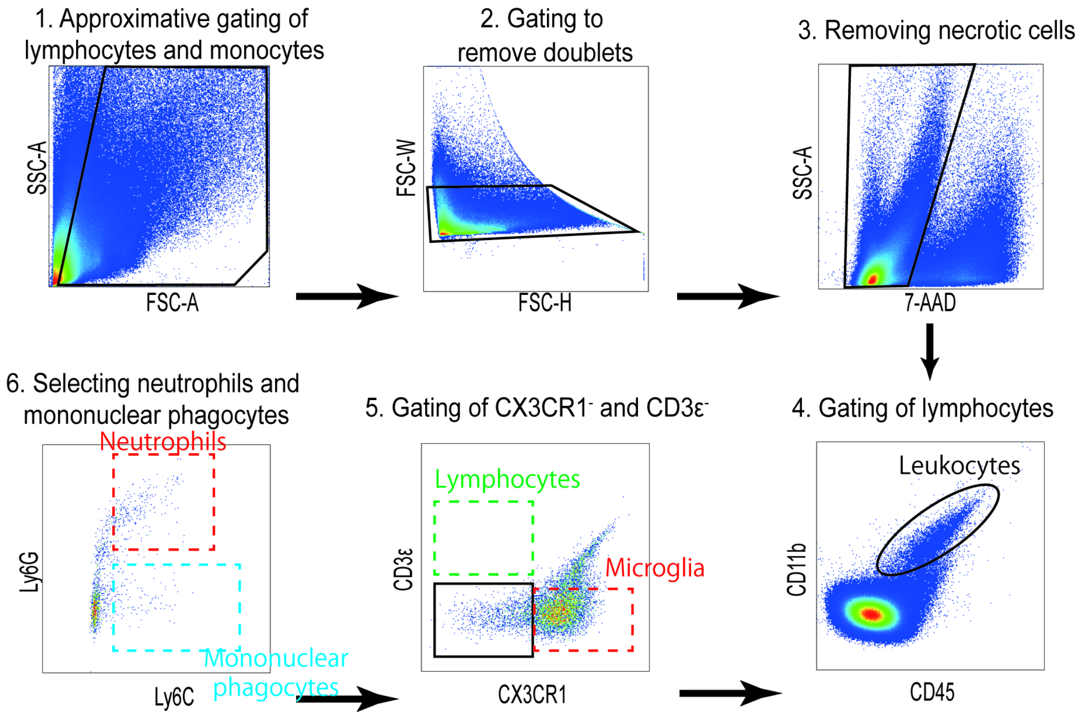


**Fig. S3.** Gating schema for immune cell populations of retinas by FACS analysis. Doublets and necrotic cells are first removed from the analysis. Immune cells are then gated as $CD45^+/CD11b^+$ (ellipse). Neutrophils are negative for microglial ($CX3CR1^+$) and lymphocyte ($CD3\varepsilon^+$) markers (rectangle). Finally, neutrophils are selected as $Ly6G^{high}$, $Ly6C^{int}$ cells (red dotted line rectangle) as opposed to monocytes, which are $Ly6G^{low}Ly6C^{high}$ (blue dotted line rectangle). Microglia: $CD45^+/CD11b^+/CX3CR1^+$, Lymphocytes: $CD45^+/CD11b^-/CD3\varepsilon^+$, Mononuclear phagocytes: $CD45^+/CD11b^+/Ly6C^{int/high}/Ly6G^{low}$, Neutrophils: $CD45^+/CD11b^+/Ly6C^{int}/Ly6G^{high}$.

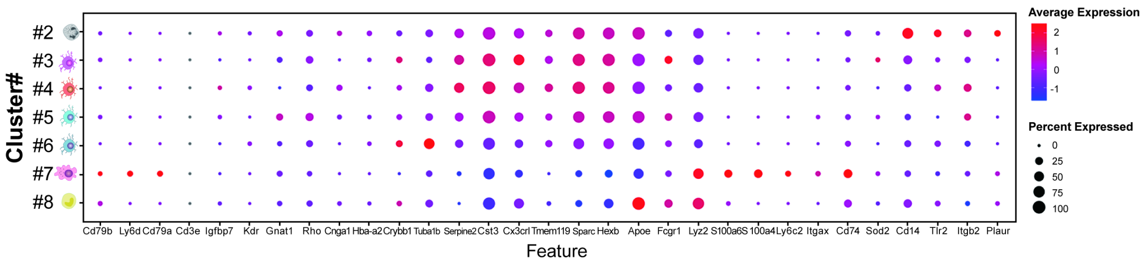


**Fig. S4.** UMAP plot of the gene activity for cell type markers. Dot plot representing expression level and frequency among cell clusters for immune cell marker genes.

Table S1. Primer sequences used for qPCR.

| Gene | Sequence | |
|---|---|---|
| *mActb* | F | 5’-GAC GGC CAG GTC ATC ACT ATT G-3’ |
| | R | 5’-CCA CAG GAT TCC ATA CCC AAG A-3’ |
| *mCcl2* | F | 5’-TAC AAG AGG ATC ACC AGC AGC-3' |
| | R | 5’-ATT CCT TCT TGG GGT CAG CAC3' |
| *mIl1b* | F | 5’-CTG GTA CAT CAG CAC CTC ACA-3’ |
| | R | 5’-GAG CTC CTT AAC ATG CCC TG-3’ |
| *mIl6* | F | 5’-AGA CAA AGC CAG AGT CCT TCA GAG A-3’ |
| | R | 5’-GCC ACT CCT TCT GTG ACT CCA GC-3’ |
| *mTnf* | F | 5’-CCC TCA CAC TCA GAT CAT CTT CT-3’ |
| | R | 5’-GCT ACG ACG TGG GCT ACA G-3’ |
